# Optimization of data-independent acquisition using predicted libraries for deep and accurate proteome profiling

**DOI:** 10.1101/2020.03.02.972570

**Authors:** Joerg Doellinger, Christian Blumenscheit, Andy Schneider, Peter Lasch

## Abstract

*In silico* spectral library prediction of all possible peptides from whole organisms has a great potential for improving proteome profiling by data-independent acquisition (DIA) and extending its scope of application. In combination with other recent improvements in the field of mass spectrometry (MS)-based proteomics, including sample preparation, peptide separation and data analysis, we aimed to uncover the full potential of such an advanced DIA strategy by optimization of the data acquisition. The results demonstrate that the combination of high-quality *in silico* libraries, reproducible and high-resolution peptide separation using micro-pillar array columns as well as neural network supported data analysis enables the use of long MS scan cycles without impairing the quantification performance. The study demonstrates that mean coefficient of variations of 4 % were obtained even at only 1.5 data points per peak (full width at half maximum) across different gradient lengths, which in turn improved proteome coverage up to more than 8000 proteins from HeLa cells using empirically-corrected libraries and more than 7000 proteins using a whole human *in silico* predicted library. These data were obtained using a Q Exactive orbitrap mass spectrometer with moderate scanning speed (12 Hz) and perform very well in comparison to recent studies using more advanced MS instruments, which underline the high potential of this optimization strategy for various applications in clinical proteomics, microbiology and molecular biology.

## INTRODUCTION

Comprehensive proteome profiling using mass spectrometry has become a key technology for studying disease mechanisms ^1^. Deep proteome coverage is most often obtained using extensive offline fractionation and data-dependent acquisition (DDA) strategies ^2,3^. The major drawbacks of this approach are the rather low throughput and the inconsistent quantification of proteins across large sample cohorts. Due to these limitations, data-independent acquisition (DIA) is an attractive alternative, as it records fragments of all peptides present in a sample and so enables consistent and accurate protein quantification ^4^. However, computational analysis of DIA data is challenging as the fragment spectra contain the information of multiple co-isolating peptides. The most successful and widespread approach is a peptide-centric data analysis strategy based on sample-specific libraries derived from DDA data ^5^. Presumably, the experimental efforts and additional instrument time needed for library generation have so far prevented an even wider use of DIA in the proteomics community. Furthermore DDA-derived libraries are biased itself due to the stochastic precursor sampling of DDA as well as the still limited sequencing speed of mass spectrometers (MS) and so contradict the unbiased nature of DIA.

Recently, *in silico* prediction of peptide spectral libraries using deep learning was introduced ^6,7^. Data-independent acquisition using *in silico* predicted libraries of whole organisms is a promising approach for deep and high-throughput proteome profiling as it eliminates experimental efforts for spectral library generation and produces unbiased high-quality libraries of all possible precursors. A major drawback of this approach is the library size which exceeds DDA-derived and sample-specific libraries by far and so impedes precursor identification. This limitation can be overcome by unbiased and sample-specific library reduction using DIA with narrow isolation windows and gas-phase fractionation (GPF) of pooled samples ^8,9^. However, this correction process of *in silico* predicted libraries requires additional instrument time just as library generation using DDA but without the need for offline peptide fractionation. Nevertheless, it is a convenient way of identifying all detectable peptides by DIA for a certain LC-MS setup in contrast to the biased DDA-derived libraries. Another current limitation of DIA is caused by the scanning speed of available MS instruments as it restricts the number of possible isolation windows for a given scan cycle duration, which allows for accurate and precise quantification. However, quantification performance of DIA is not only affected by the number of data points per peak (DPPP) provided by the MS but also depends on sample preparation, peptide separation, library generation and data analysis.

Due to recent improvements in these latter areas, including detergent-free and highly reproducible sample preparation ^10^, robust micropillar-array based peptide separation with high peak capacity ^11^, accurate *in silico* spectral library prediction ^6,7^ and sensitive and precise neural network supported data analysis ^12^, we hypothesized, that the use of longer scan cycle durations than were previously possible ^13^ could have become feasible without negatively affecting the quantification performance. This would transform into high numbers of isolation windows with decreased widths and should in turn boost precursor identification rates. Peptide co-isolation could furthermore be minimized by using isolation windows with dynamic widths based on calculations of all detectable peptides for the given LC-MS setup using empirically-corrected libraries. Based on these considerations we conducted this study with the aim to optimize isolation windows using advanced DIA strategies for improving deep and accurate proteome profiling by DIA.

## EXPERIMENTAL PROCEDURES

### Cultivation

*E. coli* K-12 (DSM 3871) and *C. albicans* strain SC5314 (ATCC MYA-2876) were cultivated on Tryptic Soy Agar (TSA) ReadyPlates^TM^ (Merck, Darmstadt, Germany) at 37°C overnight. Cells were harvested using an inoculating loop and washed in 2 × 1 mL phosphate-buffered saline (PBS) for 5 min at 4,000 × g and 4°C. HeLa cells (ATCC® CCL-2™) were cultivated in DMEM supplemented with 10 % FCS and 2 mM L–Glutamine at 37°C and harvested at 90 % confluency by scraping. Cells were washed in 2 × 2 mL PBSfor 8 min at 400 × g and 4°C.

Bacterial overnight cultures of *S. aureus* strain NCTC 8325 (DSM 4910) grown in Tryptone Soy Broth (TSB; Thermo Fisher Scientific) at 37°C were diluted to an OD600 of 0.05. 400 µl were either filled into three wells of a 6-well TPP^®^ tissue culture plate (Merck) supplemented with 0.5 % glucose and incubated under static conditions or used for inoculation of 3 × 10 mL TSB, which was incubated under rigorous mixing. After 18 h, cells from the planktonic cultivation were washed in 3 × 1 mL PBS and subsequently harvested for 5 min at 4,000 × g and 4°C. The supernatants from the biofilms were discarded and the adherent bacteria were washed using 3 × 1 mL PBS prior to mass spectrometric sample preparation.

### Sample preparation

Samples were prepared for proteomics using *Sample Preparation by Easy Extraction and Digestion (SPEED)* ^10^. At first cells were resuspended in trifluoroacetic acid (TFA) (Uvasol® for spectroscopy, Merck) (sample/TFA 1:4 (v/v)) and incubated at room temperature for 2 min (*E. coli*, HeLa,) or at 70°C for 3 min (*S. aureus*, *C. albicans*). Samples were neutralized with 2M TrisBase using 10 x volume of TFA and further incubated at 95°C for 5 min after adding Tris(2-carboxyethyl)phosphine (TCEP) to a final concentration of 10 mM and 2-Chloroacetamide (CAA) to a final concentration of 40 mM. Protein concentrations were determined by turbidity measurements at 360 nm (1 AU = 0.79 µg/µL) using GENESYS™ 10S UV-Vis Spectrophotometer (Thermo Fisher Scientific, Waltham, Massachusetts, USA), adjusted to 1 µg/µL using a 10:1 (v/v) mixture of 2M TrisBase and TFA and then diluted 1:5 with water. Digestion was carried out for 20 h at 37°C using Trypsin Gold, Mass Spectrometry Grade (Promega, Fitchburg, WI, USA) at a protein/enzyme ratio of 100:1. Resulting peptides were desalted using 200 µL StageTips packed with three Empore™ SPE Disks C18 (3M Purification, Inc., Lexington, USA) and concentrated using a vacuum concentrator ^14^. Dried peptides were suspended in 20 µL 0.1 % TFA and quantified by measuring the absorbance at 280 nm using a Nanodrop 1000 spectrophotometer (Thermo Fisher Scientific). Peptide mixtures of different species were prepared from SPEED preparations of *E. coli* K-12 and *C. albicans* as well as of a commercially available human protein digest (Promega), which was exclusively used in this experiment. The ratios of the peptide amounts of each species within the three mixtures are given in Table S1.

### Liquid chromatography and mass spectrometry

Peptides were analyzed on an EASY-nanoLC 1200 (Thermo Fisher Scientific) coupled online to a Q Exactive™ Plus mass spectrometer (Thermo Fisher Scientific). 1 µg (160 min) or 2 µg (270 min and 390 min) peptides were loaded on a μPAC™ trapping column (PharmaFluidics, Ghent, Belgium) at a flow rate of 2 µL/min for 6 min and were subsequently separated on a 200 cm μPAC™ column (PharmaFluidics) using either a 160, 270 or 390 min gradient of acetonitrile in 0.1 % formic acid at 300 nL/min flow rate (Table S2). Column temperature was kept at 50°C using a butterfly heater (Phoenix S&T, Chester, PA, USA). The Q Exactive™ Plus was operated in a data-independent (DIA) manner in the m/z range of 350 – 1,150. Full scan spectra were recorded with a resolution of 70,000 using an automatic gain control (AGC) target value of 3 × 10^6^ with a maximum injection time of 100 ms. The Full scans were followed by different numbers of DIA scans of dynamic window widths using an overlap of 0.5 Th (Table S3-S15). DIA spectra were recorded at a resolution of 17,500@200m/z using an AGC target value of 3 × 10^6^ with a maximum injection time of 55 ms and a first fixed mass of 200 Th. Normalized collision energy (NCE) was set to 25 % and default charge state was set to 3. Peptides were ionized using electrospray with a stainless steel emitter, I.D. 30 µm, (Proxeon, Odense, Denmark) at a spray voltage of 2.0 kV and a heated capillary temperature of 275°C.

### Data analysis

Protein sequences of *homo sapiens* (UP000005640, 95915 sequences, 23/5/19), *E. coli* K-12 (UP000000625, 4403 sequences, downloaded 23/5/19), *S. aureus* strain NCTC 8325 (UP000008816, 2889 sequences, downloaded 4/10/18) and *C. albicans* strain SC5314 (UP000000559, 6035 sequences, downloaded 21/11/19) were obtained from UniProt. Spectral libraries were predicted for all possible peptides with strict trypsin specificity (KR not P) in the m/z range of 350 – 1,150 with charges states of 2 – 4 and allowing up to one missed cleavage site using Prosit ^7^. Input files for library prediction were generated using EncyclopeDIA (Version 0.9.0) ^8^. The mass spectra were analyzed in DIA-NN (Version 1.6 and 1.7) using fixed mass tolerances of 10 ppm for MS^1^ and 20 ppm for MS² spectra with enabled “RT profiling” using the “robust LC” quantification strategy based on the Top3 precursors ^12^. The false discovery rate was set to 1 % for precursor identifications and proteins were grouped according to their respective genes. The resulting “report.tsv” files were filtered using R (Version 3.6) in order to keep only proteotypic peptides and proteins with protein q-values < 0.01. Peptide identifications were counted after removal of duplicated sequences resulting from the identification of multiple charge states. Proteins were quantified using the “normalised.unique” intensities. Visualization and further analysis was done in Perseus (Version 1.6.5) ^15^.

## RESULTS AND DISCUSSION

### Isolation window optimization

In an initial experiment we strived to determine the lowest number of data points per peak (dppp) necessary for accurate and precise quantification. Therefore an *in silico* generated human library (3,135,469 precursors) was empirically-corrected (137,905 precursors) based on measurements using narrow isolation windows (4 m/z widths with 2 m/z overlap) and gas-phase fractionation (8 × 100 m/z, 350 – 1,150 m/z). Afterwards, HeLa cells were measured in triplicates using a 160 min gradient in the range of 1 – 4 data points per peak in 0.5 steps. The number of isolation windows for different numbers of data points per peak was calculated from the chromatographic peak width (full width at half maximum, FWHM) of all precursors identified in the 8 gas-phase fractionations and the MS1 and DIA scan durations. The isolation window widths were calculated such as the number of precursors in the empirically-corrected library was kept constant for each window (FIGURE 1A). It was found, that the number of consistently quantified proteins increased in conjunction with the number of windows using either the whole or the empirically-corrected *in silico* predicted library (FIGURE 1B and D). However, quite strikingly, the coefficients of variations (CV) for protein quantification within the triplicate measurements did not substantially increase except when using just one data point per peak (FWHM), which led to a mean CV of 9.6 % (FIGURE 1C and E). With respect to maximize protein identifications while maintaining low CVs, the method comprising of 69 dynamic DIA windows corresponding to 1.5 data points per peak, which resulted in a mean CV of 4 %, was chosen for further evaluation of its quantification precision and accuracy using species mixtures. It should the noted, that the effective numbers of data points per peak were slightly above the calculated values, e.g. 1.8 for the 1.5 method, because the empirically-corrected library was never identified completely in the single-shot experiments.

**FIGURE 1:**
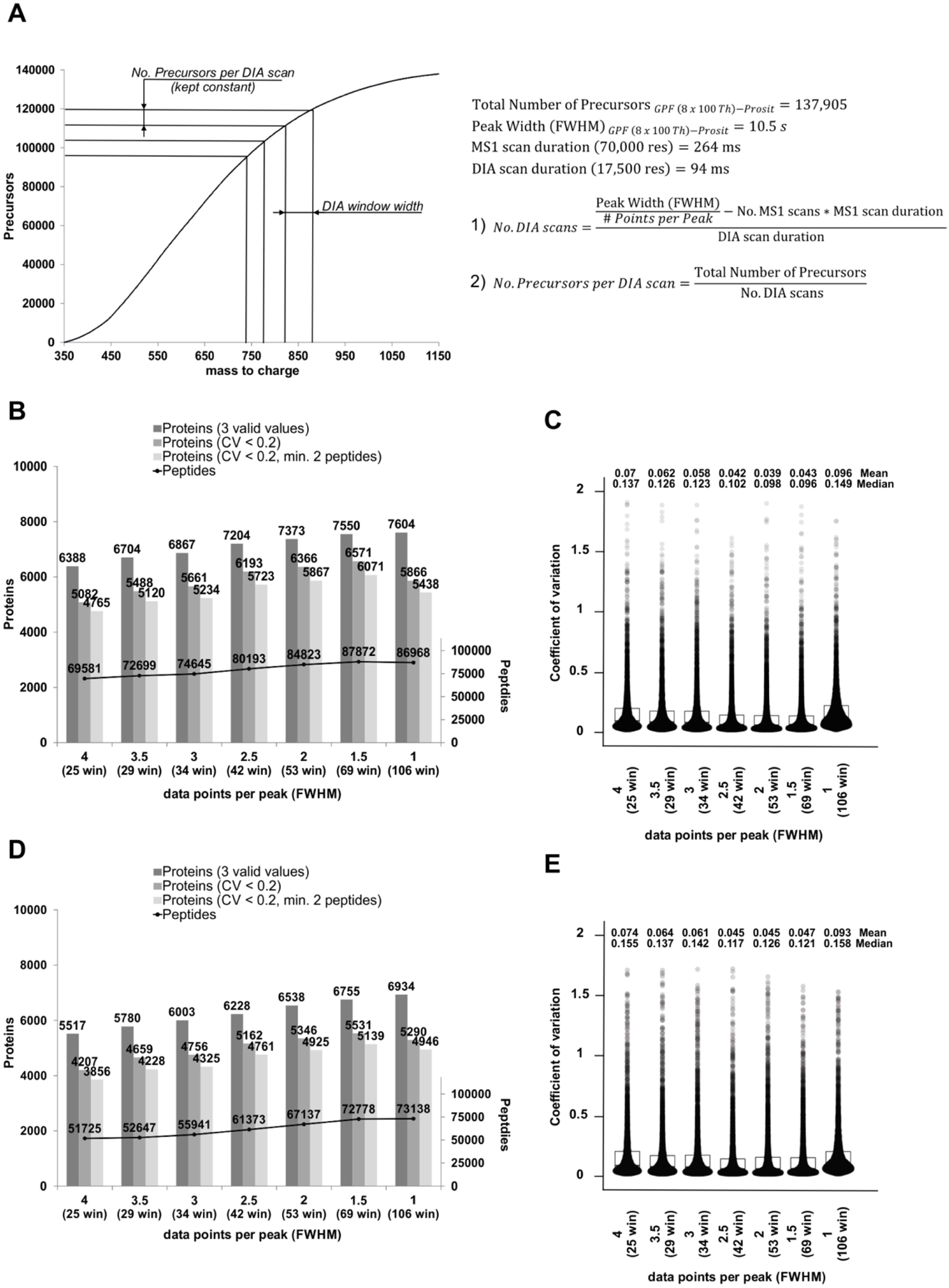
Determination of the optimum number of data points per peak. HeLa cells were analyzed in triplicates using a 160 min gradient in the range of 1 – 4 data points per peak in 0.5 steps. The number of isolation windows was calculated from the chromatographic peak width (FWHM) of all precursors identified in 8 gas-phase fractionations and the MS scan durations. The isolation window widths were calculated such as the number of precursors in the empirically-corrected library was kept constant for each window (A). Data were analyzed using either an empirically-corrected (B, C) or a whole organism *in silico* predicted library (D, E). Protein and peptide identifications using various filters (3 valid values, CV < 0.2, CV < 0.2 and minimum 2 peptides) are shown as bar plots (B, D) and the respective coefficients of variation (CV) for all proteins with 3 valid values as violin plots (C, E).

### Evaluation of quantification precision and accuracy

Quantification accuracy was analyzed using homo sapiens, *E.coli* and *C. albicans* digests, which were mixed at different ratios in order to obtain three different samples (mix 1-3) corresponding to 9 ratios ranging from −5 to 10. It was found, that accuracy of quantification varied between 0.5 – 18 % and that the actual protein ratios were in general overestimated especially for higher fold changes > 5 (FIGURE 2A). This is the exact opposite to the ratio compression issues known for label-based quantification strategies such as SILAC or TMT ^16^. Quantification precision was analyzed using two mixtures of *homo sapiens* and *E.coli* digests resulting theoretically in a two-fold protein abundance difference for the *E.coli* proteins (FIGURE 2B). T-test analysis (FDR 1 %) of 8478 quantified proteins revealed a false positive rate of 1.4 % (*homo sapiens* proteins) and a false negative rate of 7.1 % (*E.coli* proteins) considering differential abundance. This result is highly encouraging concerning the quantification precision as the actual false positive rate was highly similar to the one expected from the correction for multiple testing.

**FIGURE 2:**
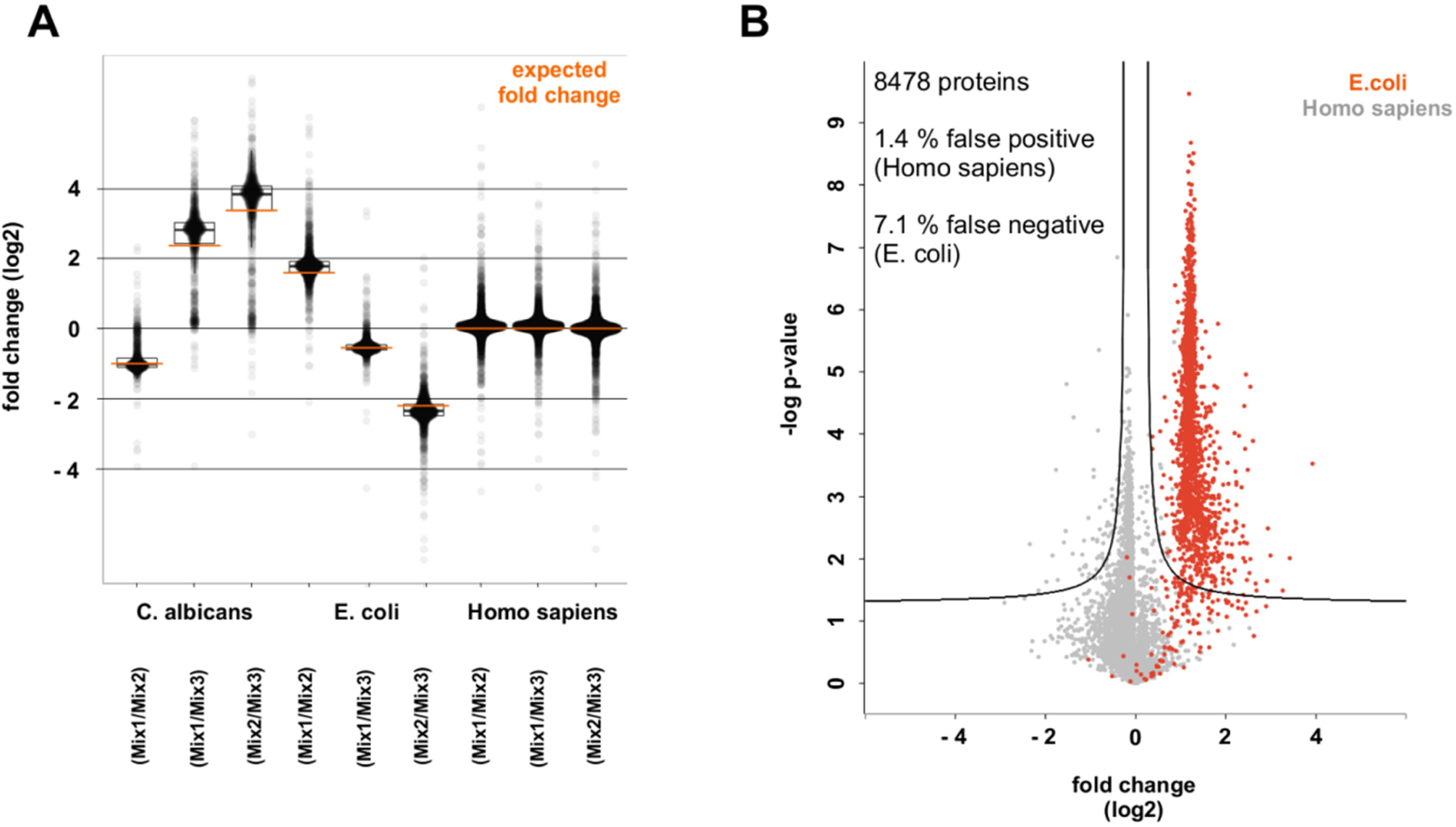
Evaluation of quantification precision and accuracy. Quantification accuracy was analyzed using *homo sapiens*, *E.coli* and *C. albicans* digests, which were mixed at different ratios in order to obtain three different samples (mix 1-3) corresponding to 9 ratios ranging from −5 to 10. The results are shown as boxplots along with the expected fold change (orange) in A. Quantification precision was studied using mixtures of *homo sapiens* and *E.coli* with an expected fold change of 2 for all *E.coli* proteins. T-test results (FDR 1 %) of 8478 quantified proteins are shown as a volcano plot in B. The false positive rate was 1.4 % (*homo sapiens*) and the false negative rate 7.1 % (*E.coli*).

### LC gradient extension for improved proteome coverage

Results from the isolation window optimization suggested that precise and accurate protein quantification is feasible even at 1.5 data points per peak (FWHM), which was further validated using species mixtures. This enables the use of high number of isolation windows, which in turn reduces peptide co-isolation and so improves peptide-centric protein identification. In order to test the current limits of this DIA approach concerning protein identifications we extended the LC gradient from 160 min to 270 min and 390 min and increased the peptide load from 1 to 2 µg for the triplicate analysis of HeLa cells. These gradients resulted in effective peptide elution times of approximately 110, 210 and 330 min (∼ 3, 5 and 7 h total analysis time) due to the large void volume of the 200 cm μPAC™ column. The isolation windows for the extended gradients were calculated based on an empirically-corrected *in silico* library exactly as described for the initial window optimization experiment. The false discovery rate (FDR) calculations were further validated by an additional analysis using combined spectra libraries of *homo sapiens* and *E.coli*. *E.coli* entries function as false targets and were used for FDR re-calculation. (TABLE 1). It was found, that Protein FDRs ranged between 0.4 – 1.0 % and peptide FDRs between 0.04 – 0.09 %, which underlines that the proposed workflow does not introduce false positive hits above the expected threshold.

**TABLE 1:** False discovery rate determination based on species mixture libraries. False discovery rate (FDR) of peptide and protein identifications were determined for HeLa analysis at different gradient lengths using a combined *in silico* predicted library of *homo sapiens* and *E.coli*.

| Sample | Points per Peak (FWHM) | Gradient [min] | Spectral Library | Peptides (Homo sapiens) | Peptides (E.coli) | Peptide Level FDR [%] | Proteins (Homo sapiens) | Proteins (E.coli) | Protein Level FDR [%] |
| --- | --- | --- | --- | --- | --- | --- | --- | --- | --- |
| HeLa | 1.5 | 160 | GPF-Prosit | 94488 | 77 | 0.08 | 8122 | 76 | 0.93 |
| HeLa | 1.5 | 160 | Prosit | 81136 | 33 | 0.04 | 7412 | 30 | 0.4 |
| HeLa | 1.5 | 270 | GPF-Prosit | 103905 | 93 | 0.09 | 8689 | 89 | 1.01 |
| HeLa | 1.5 | 270 | Prosit | 94640 | 38 | 0.04 | 8252 | 35 | 0.42 |
| HeLa | 1.5 | 390 | GPF-Prosit | 111125 | 74 | 0.07 | 8868 | 72 | 0.81 |
| HeLa | 1.5 | 390 | Prosit | 115032 | 49 | 0.04 | 8544 | 43 | 0.5 |

The gradient extension improved the number of consistently quantified proteins from 7550 (160 min) to 8234 (270 min) and 8384 (360 min) using the empirically-corrected *in silico* library and from 6755 (160 min) to 7679 (270 min) and 7879 (360 min) using the whole *in silico* library (FIGURE 3A). The moderate increase of protein identifications from 270 min to 360 min suggests that 270 min is already close to the saturation point of this LC-MS setup. The coefficients of variation were highly similar in the range of 4 – 5 % for all gradients, which underlines that the method transfer of using 1.5 data points per peak (FWHM) for extended gradients was successful (FIGURE 3B). The HeLa data for different gradient lengths were further compared to data from recent publications, which aimed to increase proteome coverage by method development ^13,17–21^ (FIGURE 3C). Although the numbers of consistently quantified proteins are not directly comparable between these publications because of different FDR calculations, protein inference strategies and number of replicates, it can be stated that the optimized DIA method performs quite well not only considering the numbers of reported proteins but also with respect to the reported mean CV values. Furthermore, the obtained data are quite remarkable when taking into account, that the Q Exactive™ Plus used in this study is an older and less advanced instrument than the ones used in the comparative studies ^13,17–21^.

**FIGURE 3:**
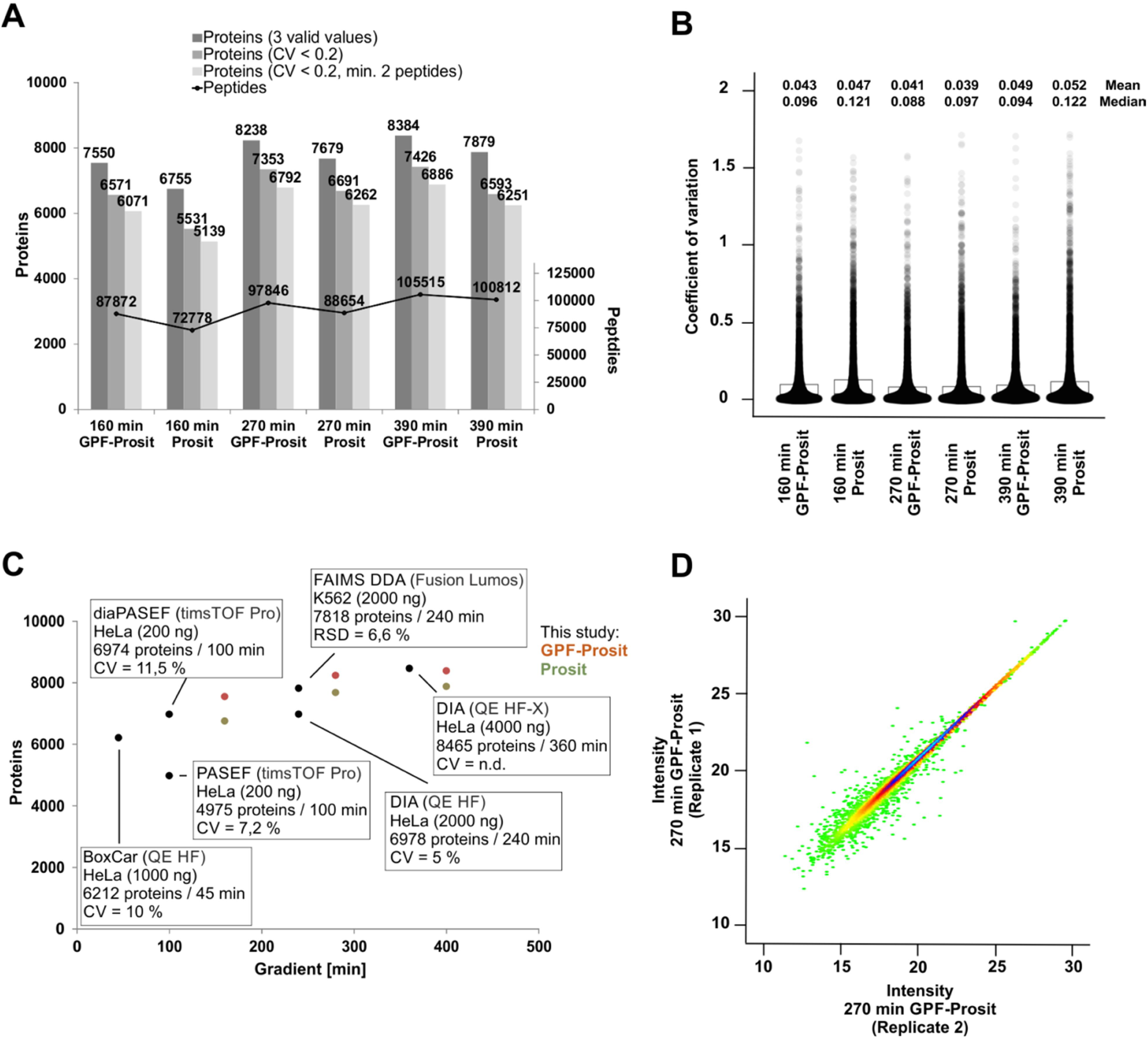
Gradient extension for improved proteome coverage. HeLa cells were measured in triplicates with LC gradients of various length (160, 270 and 390 min) using 1.5 data points per peak (FWHM). Data were analyzed using either an empirically-corrected (GPF-Prosit) or a whole organism *in silico* predicted library (Prosit). Protein and peptide identifications using various filters (3 valid values, CV < 0.2, CV < 0.2 and minimum 2 peptides) are shown as bar plots (A) and the respective coefficients of variation (CV) for all proteins with 3 valid values as violin plots (B). The results are further compared to recent publications, which aimed to increase proteome coverage by method development ^13,17–21^ (C). Correlation of the protein intensities of two replicates using a 270 min gradient is further visualized in a scatter plot (D) with the protein density distribution being color-coded.

### Application of the optimized DIA method for the analysis of *S. aureus* protein expression under different growth conditions

The ability of the optimized DIA method to gain useful insights into important biological systems was studied by analyzing *S. aureus*, which is one of the major causes of nosocomial infections worldwide ^22^. *S. aureus* pathogenesis is closely linked to glucose availability, which also enhances biofilm formation ^23^. The ability of biofilm formation increases survivability, resistance to antimicrobial substances and often virulence. It enables *S. aureus* to persist on medical devices and so facilitate spreading ^24^. Furthermore the attachment of bacteria on medical implants and host tissue is associated with chronic infections ^25^. In frame of this work, the proteome of *S. aureus* was studied at planktonic conditions as well as at elevated glucose levels under static conditions, which should induce biofilm formation and expression of virulence factors.

**FIGURE 4:**
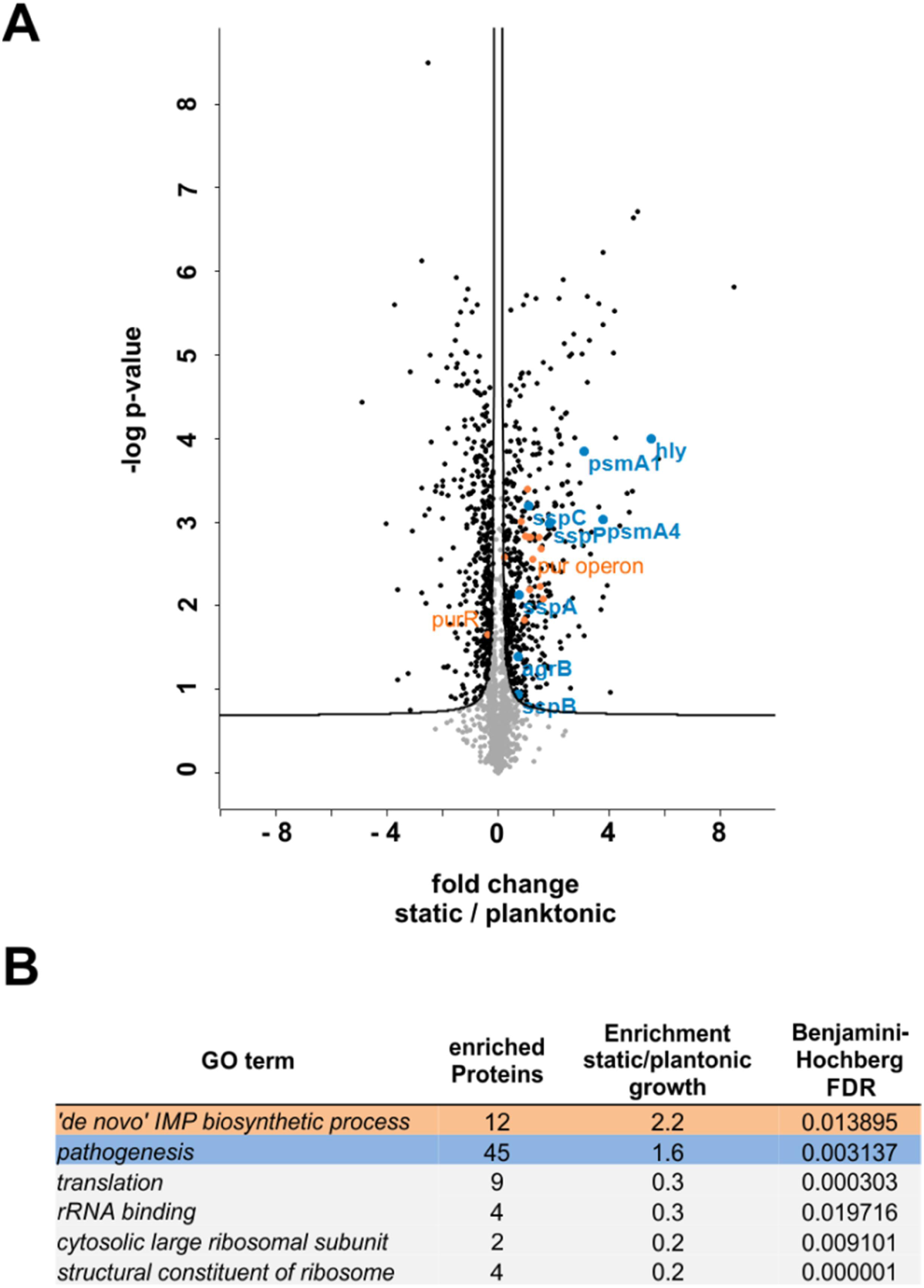
Analysis of S.aureus protein expression under different growth conditions. The proteomes of *S. aureus* grown under planktonic or static conditions with an elevated glucose level were compared. T-test results (5 % FDR) were visualized in a volcano plot (A) and revealed that the expression of 840 out of 1960 proteins was altered. Selected proteins corresponding to the GO term pathogenesis are colored blue and proteins related to purine biosynthesis, which are summarized in the GO term ‘de novo’ IMP biosynthetic process are shown in orange. The results table of the GO analysis using a Fisher-exact test (5 % FDR) is shown in B.

An *in silico* spectral library was used without experimental correction and isolation windows were not adjusted compared to the HeLa measurements. This strategy enabled the identification of 1960 proteins, which corresponds to 95 % of the number of proteins identifications present in the PeptideAtlas (http://www.peptideatlas.org). T-test analysis (5 % FDR) revealed different expression of 840 proteins between growth conditions, which are related to 7 gene ontology (GO) terms as determined using a Fisher-exact test with Benjamini-Hochberg multiple-testing correction (5 % FDR). 25 proteins related to pathogenesis [GO:0009405] were upregulated at elevated glucose levels, including alpha-hemolysin (*hly*) and several proteases, which are known to be involved in colonization and infection of human tissues (*sspA, sspB, sspC, sspP*) ^26^.

Furthermore, phenol-soluble modulins (*psmA1* and *psmA4*) were enriched under static growth conditions, which are known key biofilm structuring factors, whose expression is controlled by the quorum-sensing system (*agr*), of which *agrB* was enriched as well ^27^. The second strongly enriched GO term at elevated glucose levels was “de novo” IMP biosynthetic process [GO:0006189]. Expression of the purine biosynthesis repressor *purR* was found to be decreased, which in turn led to an increased expression of all detected genes of the *pur* operon (*purA, purB, purC, purD, purE, purF, purH, purK, purL, purM, purN, purQ, purS*). Purine biosynthesis is known to play an important role for growth and pathogenesis of *S. aureus* in human blood, which is reflected by the two essential genes *purA* and *purB* ^28^. Taken together these results demonstrate, that the optimized DIA workflow can be used for investigating complex molecular mechanisms and delivers high quality MS-data for further detailed analysis.

## CONCLUSIONS

In this study, we demonstrated that the combination of recent improvements in proteomics including sample preparation, peptide separation, spectral library generation and DIA data analysis facilitates accurate and precise proteome-wide quantification by data-independent acquisition using *in silico* predicted libraries even at very low numbers of data points per peak. The associated increase of the number of isolation windows enhanced the depth of proteome coverage for the analysis of HeLa cells to levels, which are if at all rarely described in the literature using more advanced MS instruments. The described optimization strategy could be transferred to faster scanning instruments for probably even higher proteome coverage and could be especially useful with respect to the growing interest in short gradient and high-throughput applications in clinical proteomics. Furthermore, we were able to show, that without any sample-specific optimization presumably almost complete bacterial proteomes can be analyzed directly from whole organism *in silico* predicted libraries.

## ACKNOWLEDGMENTS

*Access to proteomics data*

The mass spectrometry proteomics data have been deposited to the ProteomeXchange Consortium (http://proteomecentral.proteomexchange.org) via the PRIDE partner repository with the dataset identifiers PXD017639 () ^29^.

## CONTRIBUTIONS

J.D. and P.L. conceptualized and designed the study. J.D. and A.S. performed the experiments..

J.D. analyzed the data, prepared figures and wrote the initial draft of the manuscript. C.B. assisted with data analysis. All co-authors contributed to writing, editing, and reviewing the manuscript.

## TOC

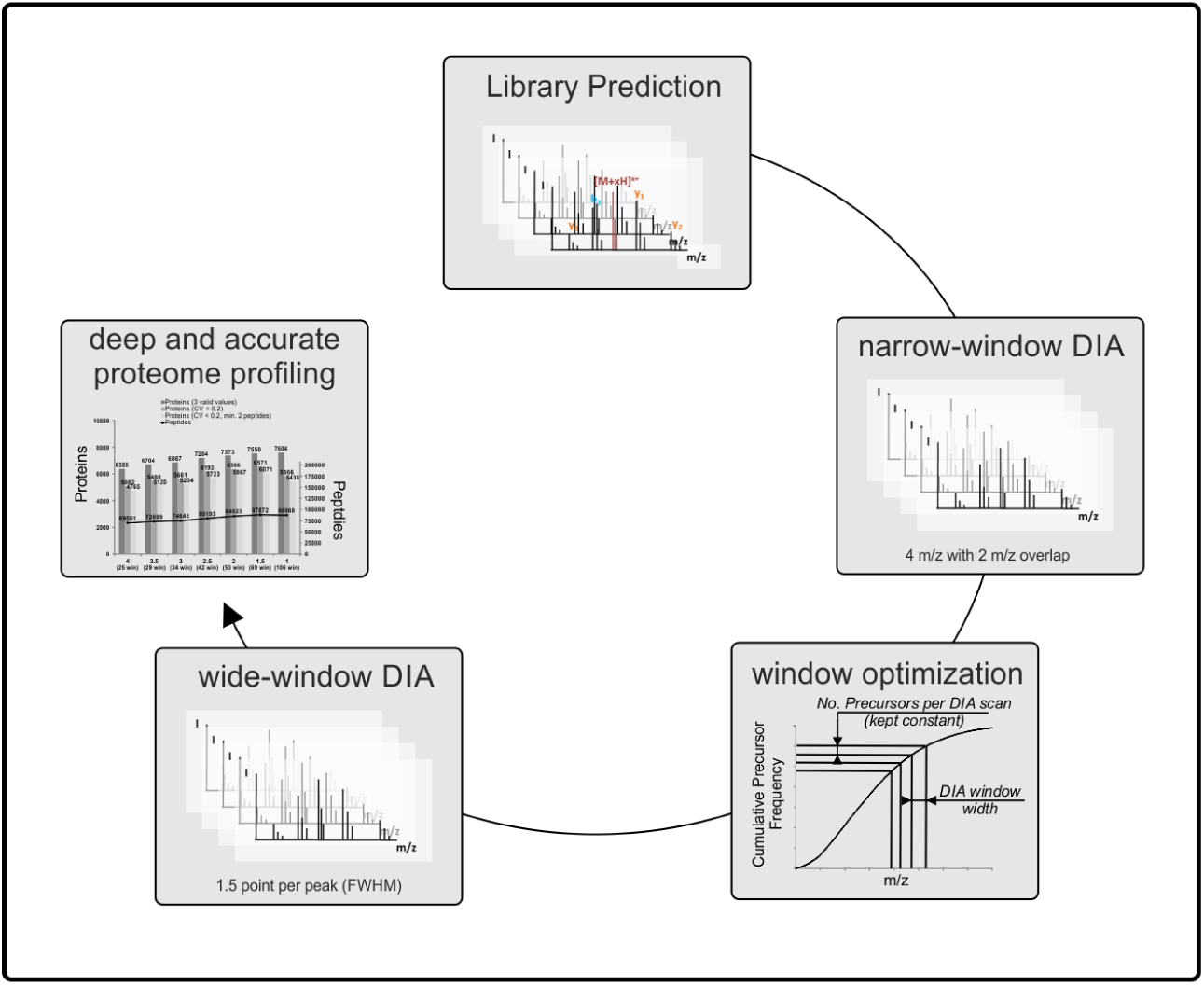

